# Structured decomposition improves systems serology prediction and interpretation

**DOI:** 10.1101/2021.01.03.425138

**Authors:** Madeleine Murphy, Scott D. Taylor, Aaron S. Meyer

## Abstract

Systems serology measurements provide a comprehensive view of humoral immunity by profiling both the antigen-binding and Fc properties of antibodies. Identifying patterns in these measurements will help to guide vaccine and therapeutic antibody development, and improve our understanding of disorders. Furthermore, consistent patterns across diseases may reflect conserved regulatory mechanisms; recognizing these may help to combine modalities such as vaccines, antibody-based interventions, and other immunotherapies to maximize protection. A common feature of systems serology studies is structured biophysical profiling across disease-relevant antigen targets, properties of antibodies’ interaction with the immune system, and serological samples. These are typically produced alongside additional measurements that are not antigen-specific. Here, we report a new form of tensor factorization, total tensor-matrix factorization (TMTF), which can greatly reduce these data into consistently observed patterns by recognizing the structure of these data. We use a previous study of HIV-infected subjects as an example. TMTF outperforms standard methods like principal components analysis in the extent of reduction possible. Data reduction, in turn, improves the prediction of immune functional responses, classification of subjects based on their HIV control status, and interpretation of these resulting models. Interpretability is improved specifically by applying further data reduction, separation of the Fc from antigen-binding effects, and recognizing consistent patterns across individual measurements. Therefore, we propose that TMTF will be an effective general strategy for exploring and using systems serology.

**Summary points:**

- Structured decomposition provides substantial data reduction without loss of information.
- Predictions based on decomposed factors are accurate and robust to missing measurements.
- Decomposition structure improves the interpretability of modeling results.
- Decomposed factors represent meaningful patterns in the HIV humoral response.

## Introduction

Whether during a natural infection, therapeutic vaccination, or an exogenously administered antibody therapy, antibody-mediated protection is a central component of the immune system. While the unique property of antibodies is conceptually simple—they undergo affinity enrichment toward specific antigens—the mechanisms of resulting protection are mediated through a network of interactions.^1^ Therapies are often optimized based upon the titer or neutralizing capacity of the antibodies they deliver. However, many of the mechanisms for antibody-mediated protection occur through secondary interactions with the immune system via an antibody’s fragment-crystallizable (Fc) region. While more challenging to quantify and identify as the mechanism of protective immunity, these immune system responses, such as antibody-dependent cellular cytotoxicity (ADCC),^2,3^ complement deposition (ADCD),^4^ cellular phagocytosis,^5^ and respiratory burst^6^ are known to be just as or more important in many diseases.

A suite of recent technologies promises to broaden our view of antibody-mediated protection as the microarray did for gene expression. Systems serology aims to broadly profile the humoral immune response by quantifying both the antigen-binding and immune interaction of antibodies in parallel.^7^ In these assays, antibodies are first separated based on their binding to a panel of disease-relevant antigens.^8^ Next, the binding of those immobilized antibodies to a panel of immune receptors is quantified. Other molecular properties of the disease-specific antibody fraction that affect immune engagement, such as glycosylation, may be quantified in parallel in an antigen-specific or -generic manner.^8,9^ By accounting for the two necessary events for effector response—antigen binding and immune receptor engagement—these measurements have proven to be highly predictive of effector cell-elicited responses and overall antibody-elicited immune protection.^10^

Though systems serology provides a major advancement in our ability to analyze the antibody-elicited immune response, analysis of these data is in a nascent stage. Standard machine learning methods, such as regularized regression, principal components analysis, and partial least squares regression have been effective in identifying highly predictive immune correlates of protection.^11,12^ However, identifying how specific molecular changes give rise to protection remains challenging. First, because many of the measurements are overlapping in the molecules they quantify, or measure co-dependent processes, much of the data is highly inter-correlated.^13,14^ Particularly when analyzing polyclonal antibody responses such as those which arise in vaccination or natural infection, protection may arise through single or combinations of molecular species and features within the antibody response, through either individual or combinations of antigens.^15,16^ Ultimately, improvements in our ability to identify patterns in these data, and how they relate to protection, should span the contribution of individual molecular changes to overall immune protection with mechanistic detail. To date, no methods have provided a means to holistically visualize the variation in serology measurements.

While systems serology measurements include a variety of different assays to quantify humoral response, a common overall structure exists to the data. The vast majority of the measurements quantify the extent to which an antibody bridges all pairs of target antigen and receptor panels, across a set of individuals.^7^ These measurements, therefore, can be thought of as a three-dimensional dataset, where every number in this cube of data represents a single measurement (Fig. 1A/B). Then, separately from these measurements, some antigen-generic properties of the humoral response, such as overall antibody glycosylation, may be assessed.^9,17^ With data of three or more dimensions, a family of methods called tensor factorization provides a generalization of matrix decomposition methods like principal components analysis (PCA).^18^ These methods are especially effective at data reduction when measurements have meaningful multi-dimensional features, such as time-course measurements.^19^ Like PCA, tensor decomposition methods, when appropriately matched to the structure of data, help to visualize its variation, reduce noise, impute missing values, and reduce dimensionality.^20^

**Figure 1:**
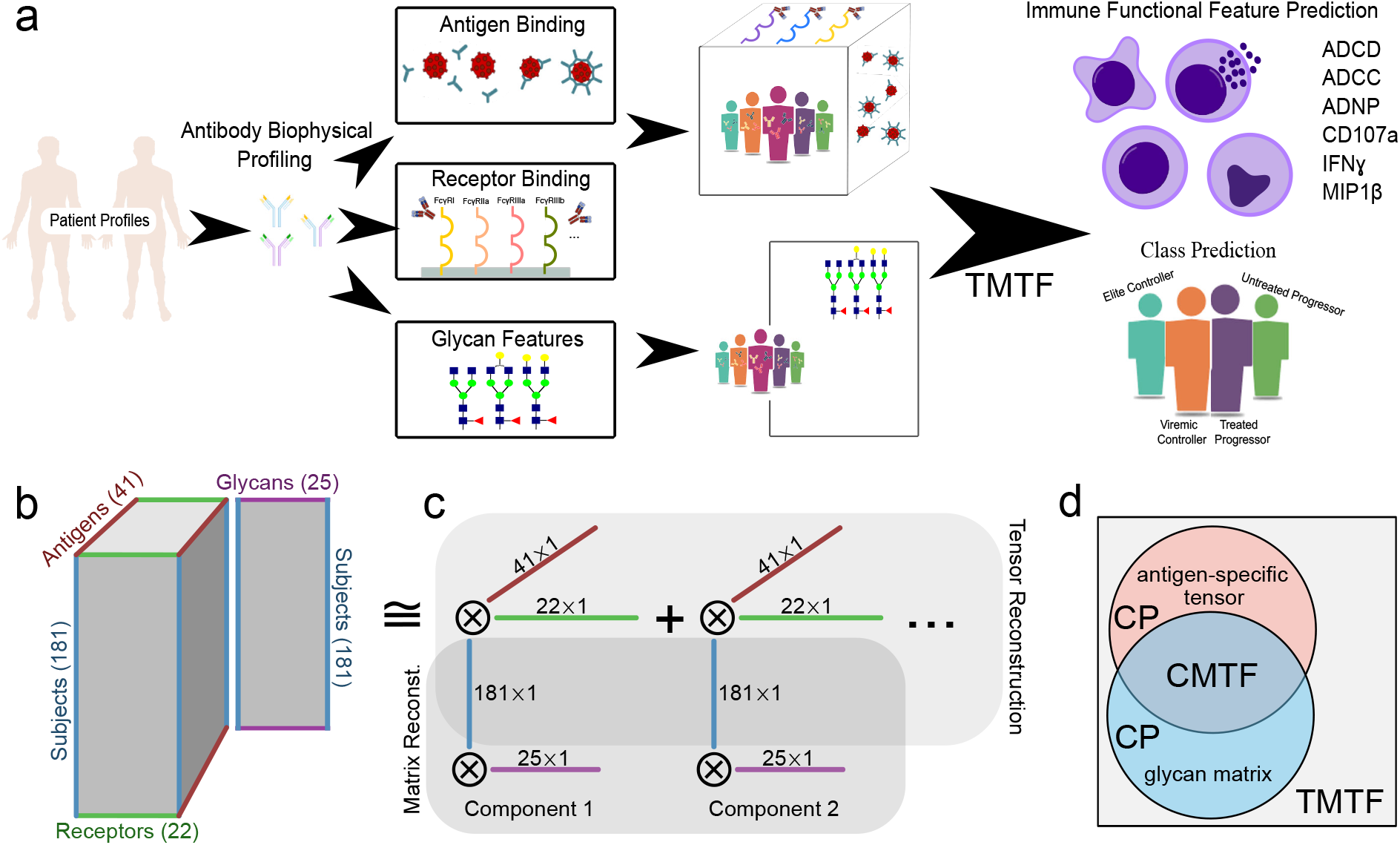
Systems serology measurements have a consistent multi-dimensional structure. A) General description of the data. Antibodies are first separated based on their binding to a panel of disease-relevant antigens. Next, the binding of those immobilized antibodies to a panel of immune receptors is quantified. Other molecular properties of the disease-specific antibody fraction that affect immune engagement, such as glycosylation, may be quantified in parallel in an antigen-specific or - generic manner. These measurements have been shown to predict both disease status (see methods) and immune functional properties—ADCD, ADCC, antibody-dependent neutrophil phagocytosis (ADNP), and natural killer cell activation measured by IFNγ, CD107a, and MIP1β expression. B) Overall structure of the data. Antigen-specific measurements can be arranged in a three-dimensional tensor wherein one dimension each indicates subject, antigen, and receptor. In parallel, antigen-generic measurements such as quantification of glycan composition can be arranged in a matrix with each subject along one dimension, and each glycan feature along the other. While the tensor and matrix differ in their dimensionality, they share a common subject dimension. C) The data is reduced by identifying additively-separable components represented by the outer product of vectors along each dimension. The subjects dimension is shared across both the tensor and matrix reconstruction. D) Venn diagram of the variance explained by each factorization method. Canonical polyadic (CP) decomposition can explain the variation present within either the antigen-specific tensor or glycan matrix on their own.^20^ CMTF allows one to explain the shared variation between the matrix and tensor.^21^ In contrast, here we wish to explain the total variation across both the tensor and matrix. This is accomplished with TMTF (see methods).

In this work, we use the structure of systems serology measurements to improve its visualization, infer missing values, and predict functional immune features. As an example, we analyze a study wherein systems serology measurements were shown to predict both functional immune responses and disease status within HIV-infected subjects.^12^ We first develop a new tensor factorization approach— total matrix-tensor factorization (TMTF)—to decompose these measurements into consistent patterns across subjects, immunologic features, and antigen targets. Inspecting these factors reveals interpretable patterns in the humoral response, and these pattern’s abundance across subjects predicts functional immune responses and subjects’ HIV infection state. Importantly, TMTF significantly improves the interpretability of these predictions compared to methods that do not recognize the structure of these data. This approach, therefore, provides a very general data-driven strategy for improving systems serology analysis.

## Results

### Systems serology measurements can be drastically reduced without loss of information

We first sought to determine whether the structure of systems serology measurements could inform better data reductio strategies (Fig. 1). To integrate antigen-specific and -generic measurements, we developed a new form of tensor decomposition we will refer to as total matrix-tensor factorization (TMTF) (Fig. 1C). By concatenating both the unfolded tensor and matrix during the alternating least squares (ALS) solve for the subject dimension, we maximize the variance explained across both datasets (Fig. 1D, see methods). This is in contrast to coupled matrix-tensor factorization, which only explains the variance shared between the datasets^21^ (Fig. 1D).

To determine the extent of data reduction possible, we examined the reconstruction error upon decomposition with varying numbers of components (Fig. 2A). As we start with a 181×22×41 tensor and 181×25 matrix, we start with a dataset of 168,000 values, of which 96,000 or 57% were measured and therefore not missing. After factorization with 10 components, we are left with four matrices of 181×10, 22×10, 41×10, and 25×10. Therefore, we reduce the dataset to 2.8% of the size (2,690 numbers), while preserving 93% of its variation (Fig. 2A). We compared this to the dimensionality reduction possible with PCA and the data organized in a “attened matrix form (Fig. 2B). TMTF consistently led to a similar accuracy on reconstruction with half the resulting factorization size compared to PCA (Fig. 2B). For example, TMTF led to a normalized unexplained variance of 0.15 at ∼1,024 values within the factorization, while PCA required ∼2,048 to do the same. This gave us confidence that this structured factorization can greatly reduce the data while preserving its meaningful variation.

**Figure 2:**
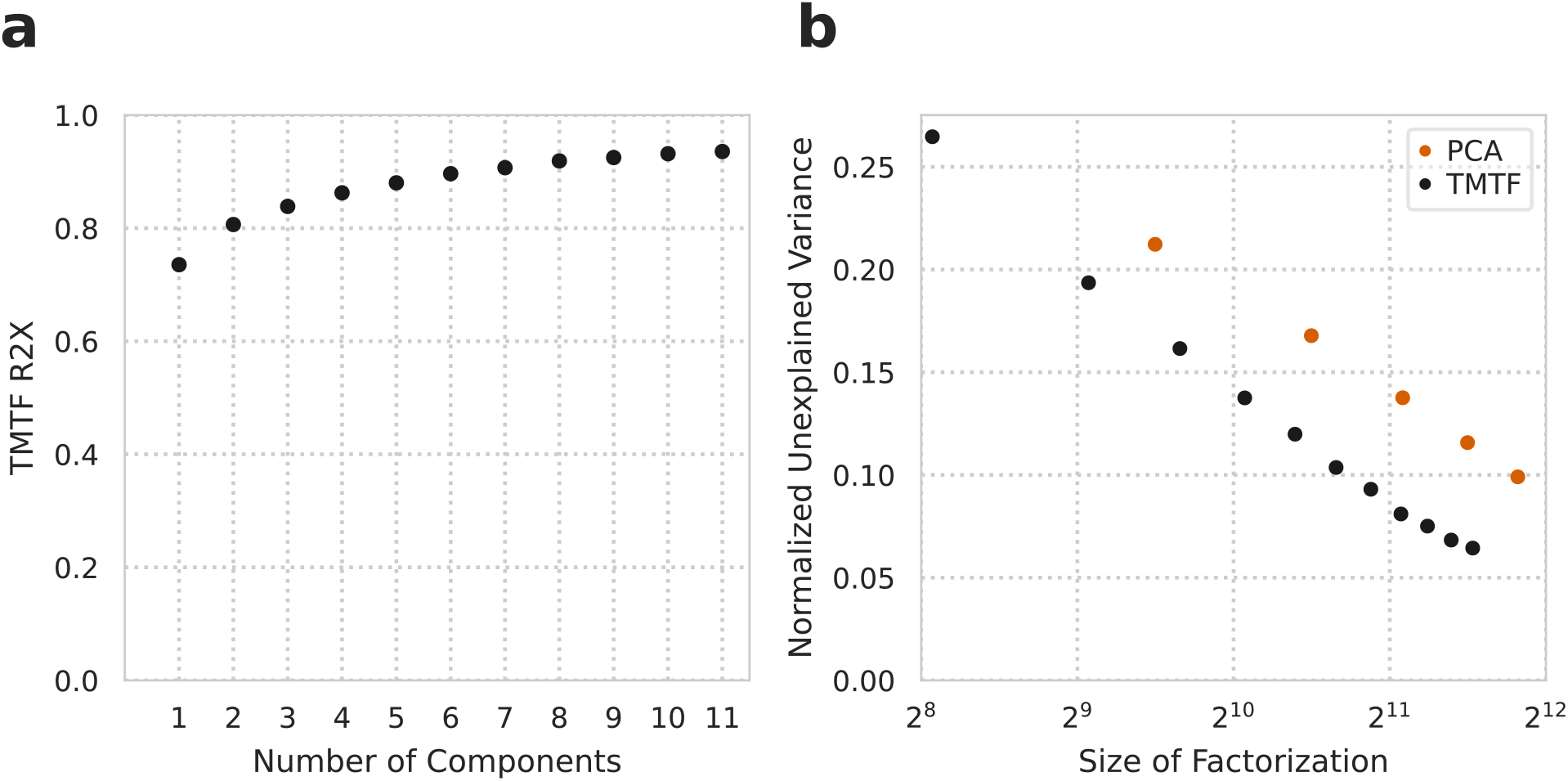
TMTF improves data reduction of systems serology measurements. A) Percent variance reconstructed (R2X) versus the number of components used in TMTF decomposition. B) TMTF reconstruction error compared to PCA over varying sizes of the resulting factorization. The unexplained variance is normalized to the starting variance. Note the log scale on the x-axis.

### Factorization accurately imputes missing values

With confidence that factorization was identifying consistent patterns within the HIV serology response, we wondered if our approach might improve missing values imputation. Systems serology measurements require a panoply of different measurements in combination; especially in such large-scale efforts, measurements can be missing for a variety of reasons. Subject samples can be limited, and so only be available for a small set of measurements. A subset of measurements may be of particular interest to investigators; for example, surface antigens to a virus may be prioritized for more detailed Fcγ receptor binding measurements, since they are more likely to be exposed in the extracellular space. In our example HIV serology data, many antigen-receptor pairs are absent, as are glycan measurements for half of the subjects. In such a case traditional regression models were limited to a smaller subset of either the data or the subjects in making predictions.^12^ TMTF provides the possibility of eliminating this tradeoff while simultaneously imputing the absent data. Effective imputation also enables outlier detection to, for example, identify assays that may have technical problems. Finally, observing that factorization accurately imputes missing values further supports that this approach identifies biologically-meaningful and consistent patterns.

To evaluate whether factorization could accurately impute missing values, we artificially introduced missing values by randomly removing *entire* receptor-antigen pairs across all subjects. TMTF was then performed which effectively filled these in, and we then calculated the variance inferred in this left-out data (Fig. 3A). Factorization imputed these values with similar accuracy to the variance explained within observed measurements (Fig. 2A), supporting that it is able to identify meaningful patterns even in the presence of missing measurements. This provides additional evidence that the patterns identified through factorization are a meaningful and substantial representation of the data.

**Figure 3:**
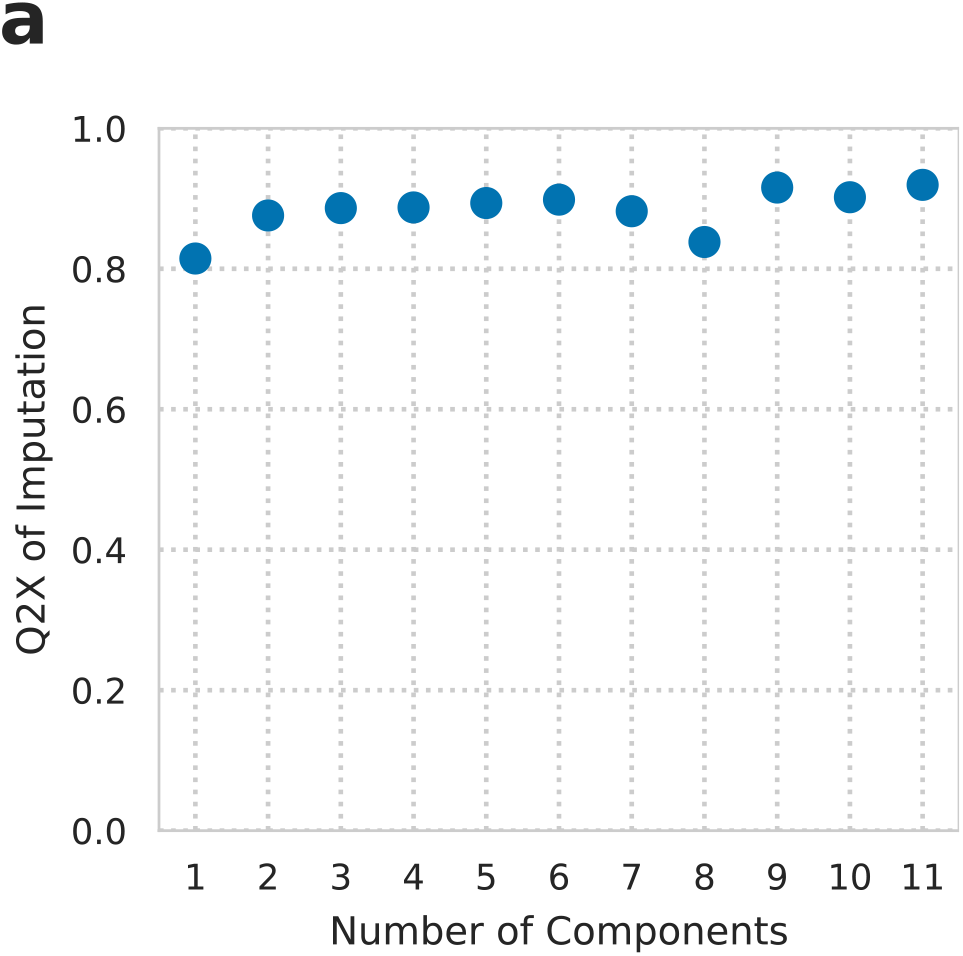
TMTF accurately imputes missing values. Percent variance predicted (Q2X) versus the number of components used for imputation of held out receptor-antigen pairs.

### Factor components represent consistent patterns in the HIV humoral immune response

Systems serology measurements have typically been analyzed by standard regularized prediction methods, such as elastic net and partial least squares regression, on the measurements themselves.^12,22^ These methods are very effective at prediction and do provide some interpretation of the mechanism behind those predictions. At the same time, a per-measurement perspective can hide that immune responses occur through polyclonal responses across antigens and through regulatory changes that affect multiple receptors. Systems serology measurements further complicate interpretation due to the combined influence of both antigen binding and immune receptor interaction differences.^23^

In contrast, tensor factorization provides a much more interpretable view of the coordinate changes along these dimensions (Fig. 4). To visualize the resulting factors, we plotted all of the components along their subject (Fig. 4A), receptor (Fig. 4B), antigen (Fig. 4C), and glycan (Fig. 4D) factors. Notably, these factors quantitatively visualize these measurements in their totality, which has not been possible to date. To interpret the resulting plots, one can examine the values of an individual component along each plot, as each component represents a separate pattern of variation within the data (additively separable variation), and each plot represents a separate dimension. The effect of each component is the product of its factors along each dimension, and so we can examine the contribution of each dimension individually.

**Figure 4:**
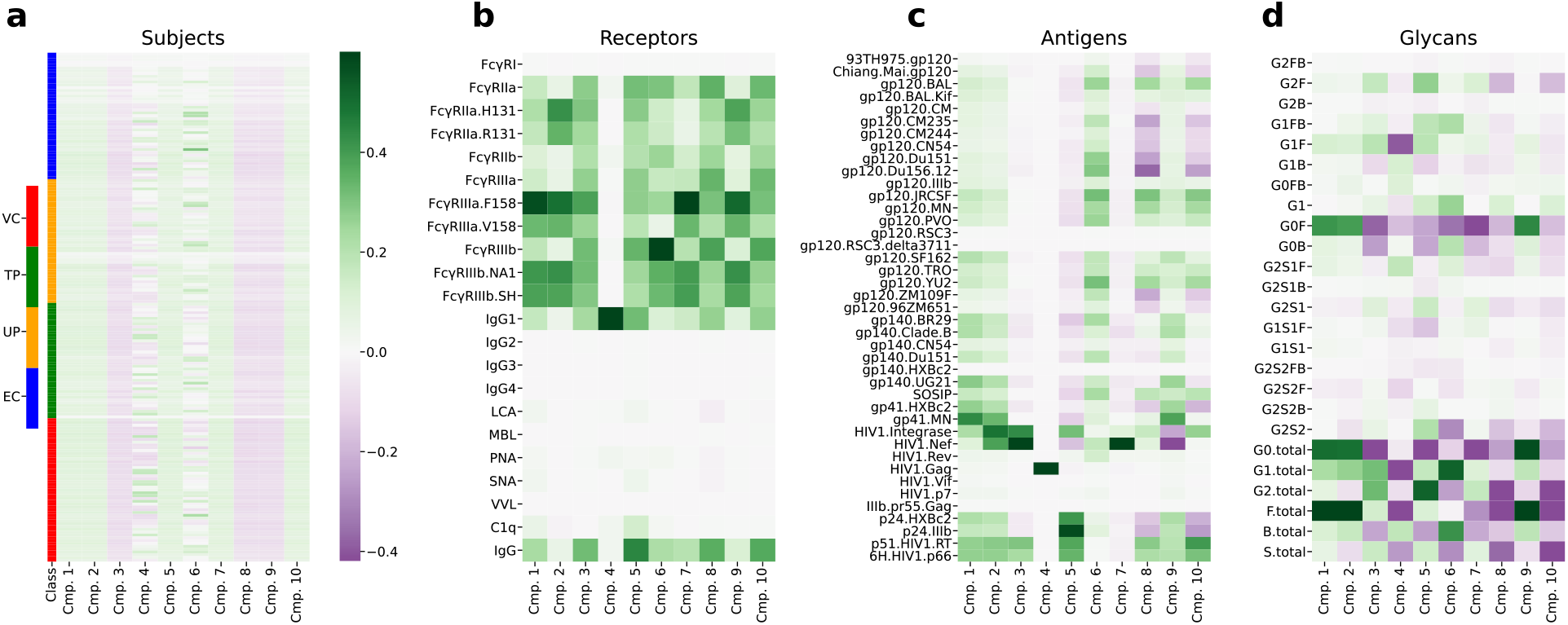
Factor components represent consistent patterns in the HIV humoral immune response. Decomposed components along subjects (A), receptors (B), antigens (C), and glycans (D). EC: Elite Controller, TP: Treated Progressor, UP: Untreated Progressor, VC: Viremic Controller (see methods). All plots are shown on a common color scale. Measurements were not normalized, and so magnitudes within a component are meaningful. Antigen names indicate both the protein (e.g., gp120, gp140, gp41, Nef, Gag) and strain (e.g., Mai, BR29).

Through inspection of each individual component, one can understand the consistent patterns across the serology measurements. For instance, component 4 explains a pattern of variation specifically in IgG1 (Fig. 4B) and Gag (Fig. 4C). The amount of this Gag-binding IgG1 varies according to each subject’s component 4 value (Fig. 4A). Finally, variation along this component corresponds to variation in glycan composition, and most prominently a decrease in G1F and G1 overall (Fig. 4D). As each component is additively independent, they can be inspected on a component-by-component basis to explore every pattern of variation within the serology.

The factorization revealed many interesting quantitative patterns. For example, component 5 represents a shift between surface antigen (gp120, gp140) and p24/p51/p66 binding (Fig. 4C), which would be nearly impossible to identify without separating receptor and antigen effects as TMTF does. We identified that some technical artifact resulted in genotype-specific FcγRIIIa measurements that are more sensitive than the genotype-generic ones (Fig. S2). Both components that we interpreted as explaining high-affinity FcγRIIIa and FcγRIIIb binding due to preferential weighting of the less sensitive measurement (Fig. 4B)—6, 8, and 10—also had strongly negative fucosylation and G0F weights (Fig. 4D). These glycosylation changes are known to hinder FcγRIIIa interaction and therefore ADCC.^24^ C1q binding was most heavily weighted along component 5 (Fig. 4B), which also weighted coordinate galactosylation and fucosylation (Fig. 4D, G2F/G1FB/G1F), which are known to promote this binding.^24^ These observations support that the TMTF components represent meaningful variation among both the antigen-specific and glycosylation measurements, and that they can be used to clearly derive mechanistic insight.

### Structured data decomposition accurately predicts functional measurements and subject classes

Next, we evaluated whether our reduced factors could predict the functional responses of immune cells and subject classes. We attempted to predict previous functional data for antibody-dependent complement deposition (ADCD); cellular cytotoxicity (ADCC); neutrophil phagocytosis (ADNP); and level of natural killer cell activation determined by expression of IFNγ, CD107a, and MIP1β. Separately, we predicted broad subject disease statuses: Controller versus Progressor, and Viremic vs Non-Viremic (see methods). As a baseline of comparison, we reimplemented the immune functionality and subject predictions previously applied to these data.^12^ We observed similar performance to that reported (Fig. 5). Differences from earlier results could be explained by corrections we made to the cross-validation strategy to prevent over-fitting (see methods).

**Figure 5:**
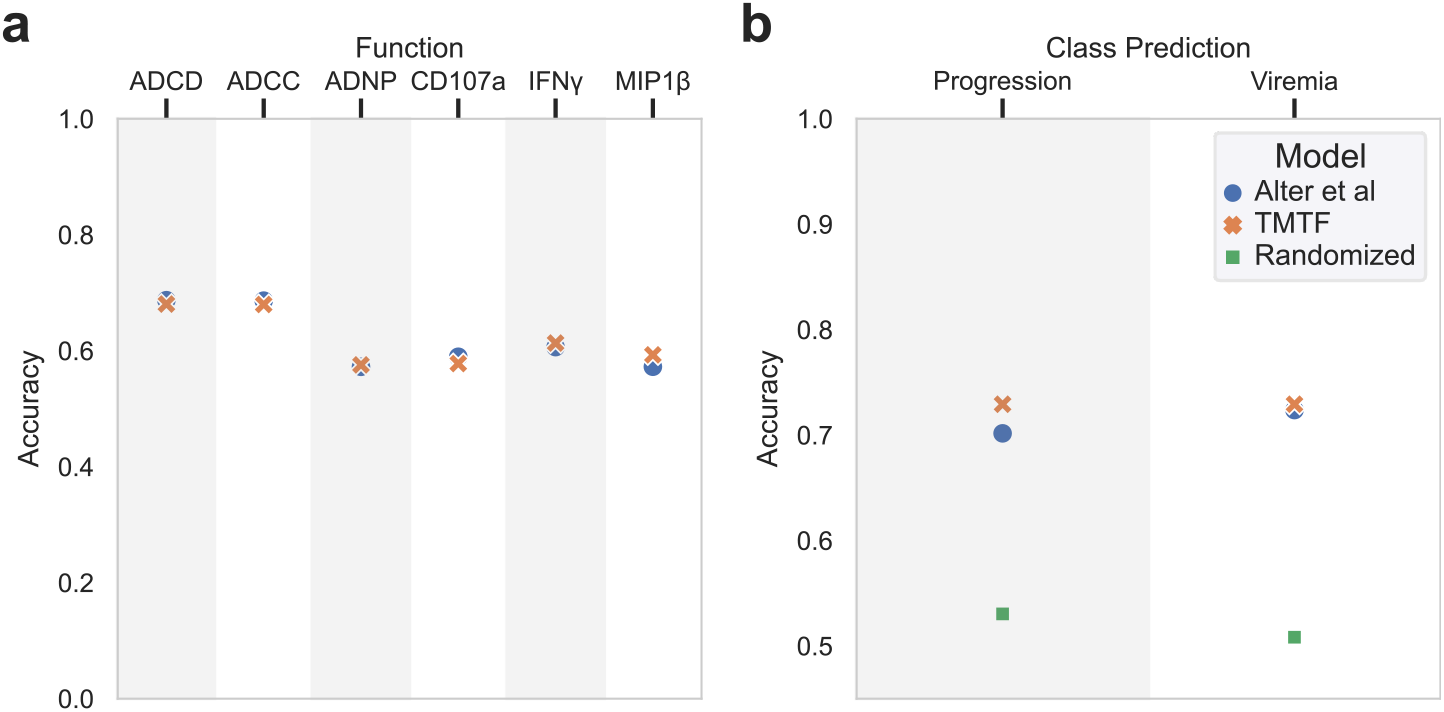
Structured data decomposition more accurately predicts functional measurements and subject classes. (A) Accuracy of prediction (defined as the Pearson correlation coefficient) for different functional response measurements. (B) Prediction accuracy for subject viral and controller status.

We benchmarked the ability of the factorized results to make these same predictions. TMTF represents the variation among subjects within a subject matrix (Fig. 4A). To make predictions, one needs only work with these subject factors rather than the hundreds of original variables in the entire dataset. Furthermore, TMTF provides factorization results for each subject despite the presence of missing values. We therefore were able integrate the glycan data with the other serology measurements, allowing us to utilize all data gathered and make observations across the datasets.

Particularly as we observed systematic, non-linear trends between the genotype-specific and -generic measurements across antigens (Fig. S2), we surmised that there might be a non-linear relationship between our factors and each prediction. Greatly reducing the number of variables to 10 components allowed us to apply a Gaussian process model to predict each function and subject class, allowing for non-linear effects.^25^ We separated our quantification of model accuracy based on whether a subject was included or excluded previously.^12^

Broadly, we saw nearly identical performance in predicting immune functional responses (Fig. 5A). This was also the case for predicting whether subjects were classified as progressors or viremic (Fig. 5B). Therefore, we concluded that TMTF preserves sufficient information to predict these important features.

### Immune function correlates match biological mechanisms

Inspecting our predictive models revealed mechanistic insights that correspond to the known biology of each immune functional response. ADCD was predicted almost exclusively using component 5 (Fig. 6A). Component 5 carried the strongest weight among components for C1q binding and so likely indicates classical pathway activation.^26^ It also showed positive weighting of LCA, PNA, and SNA binding, indicative of lectin-pathway complement activation (Fig. 4B).^26^ The regression model in the original study also identified C1q binding as important,^12^ but was unable to capture the relationship to the lectins or the shift between surface antigen and p24/p51/p66 binding (Fig. 4C). This further insight is made possible by the ability of TMTF to separate out individual factors in the serology data.

**Figure 6:**
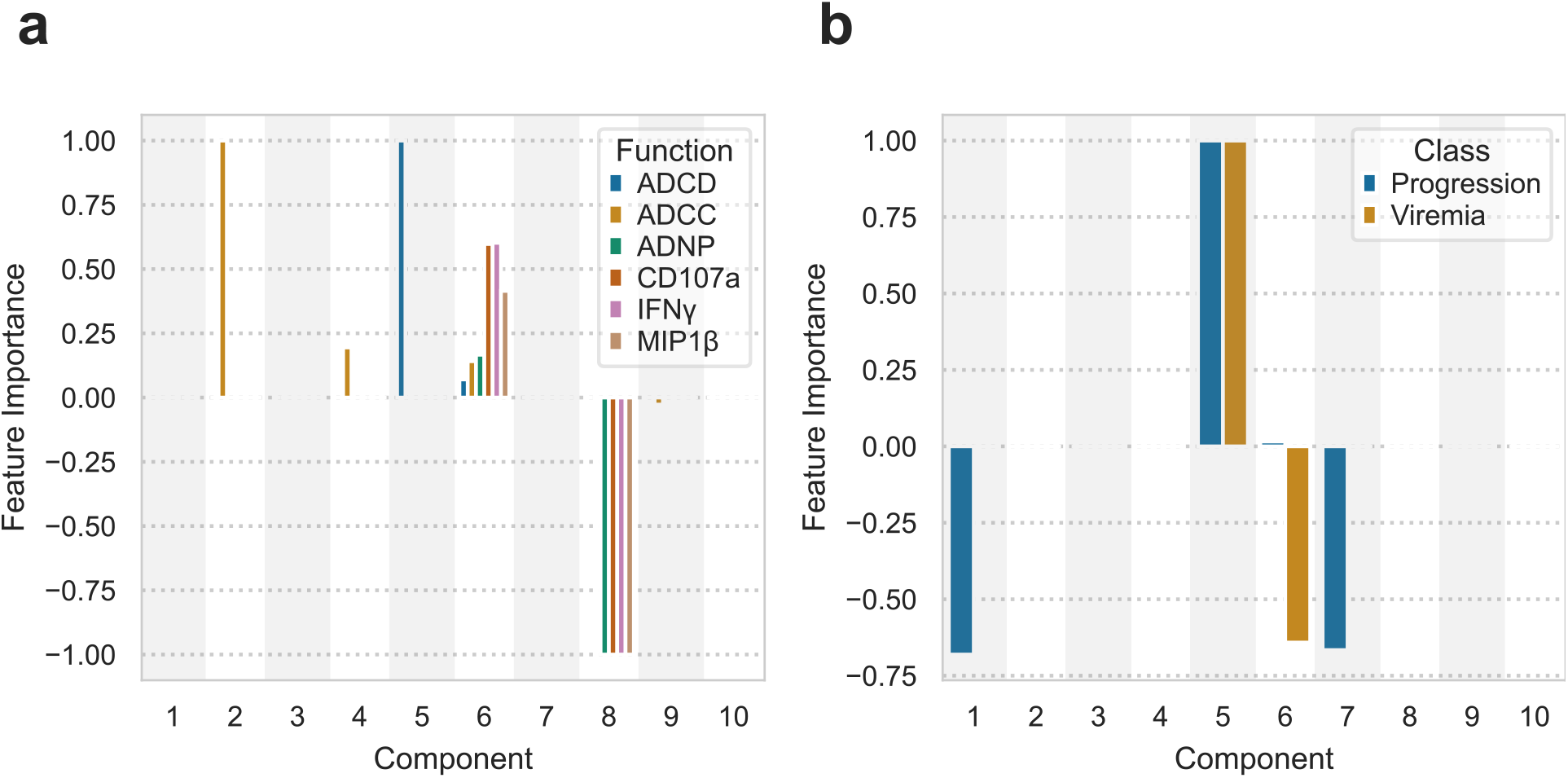
Model component effects. Model component effects for each function (A) and subject class (B) prediction. Component effects are quantified using the variable weights for a linear model, and the inverse RBF kernel length scale for a Gaussian process model.^27^ For the Gaussian process component effects, the component effect is also multiplied by the sign of the corresponding linear model to show whether that variable has an overall positive or negative effect. The component effects are shown scaled to the largest magnitude within each model.

Similarly, our model provides new, clearer observations of relationships between the factorized results and ADCC. ADCC was most prominently predicted through component 2, and secondarily 4 and 6 (Fig. 6A). Component 2 represents broad antigen binding across both gp120 and gp140, along with differences between the genotype-specific and -generic receptor measurements (Fig. 4B/C). As discussed above, we believe this is due to a difference in sensitivity of the two assays. Broadly, the receptor profile seems to only weight the more sensitive measurements (Fig. S2). We believe this component represents greater antigen binding overall, picked up by even the less sensitive assays, which also explains the positive correlation with all glycoforms (Fig. 4D). Component 6 again has broad binding across surface antigens but weights the genotype-generic FcγRIIIa and FcγRIIIb measurements more—we expect this represents high-affinity or high-avidity surface binding.

Antibody-dependent neutrophil-mediated phagocytosis (ADNP) was most prominently explained by components 6, and (negatively) 8 (Fig. 6A). These highlight two mechanisms of driving neutrophil phagocytosis—via lectin engagement (components 6 and 8) and/or FcγR binding (component 6). As described above, we interpret component 6 to indicate high-affinity or high-avidity receptor binding across surface antigens.

All three cytokines, used as measures of NK cell activation, only depended upon component 6 and (negatively) 8 (Fig. 6A). We should expect similar models across these cytokines as their amounts are highly correlated across subjects and were all measured as surrogates of NK cell activation.^12^ The greater dependence on component 6 compared to ADCC is consistent with our expectations for the determinants of each signal; ADCC might be driven by high amounts of FcγRIIIa engagement from a mix of high- and low-affinity/avidity interactions, while NK cell activation provides a quantitative activation measurement that is best promoted through exclusively high-affinity/avidity engagement. Overall, our model interpretation supports that the TMTF-decomposed factors identify patterns that are relevant to immune functional responses.

### HIV progression and viremia are predicted by specific, multi-antigen responses

The clinical properties of a disease likely involve coordinated immune functions. As a result, being able to reason about a small number of consistent immunological patterns might be especially helpful for interpreting predictive models. Therefore, we took a similar approach for evaluating the variable dependence for predictions of subject progression and viremic status. Remarkably, progression was exclusively explained by just components 5 and, to a lesser degree, 1 and 7 (Fig. 6B). In fact, a model using only component 5 was just as predictive as one with all components available. Component 5 corresponds to increased amounts of p24 binding across strains. Abundance of p24 antigen and its antibody titer has been proposed as an effective marker of HIV progression,^28^ and predictive of death,^29^ although much of this is via correlation with viral RNA and CD4+ counts.^30^ This component also displays negative weighting across gp120 and gp140 antigens, likely reflecting a decrease in antibody titers overall, which has separately been found to predict progression.^31^ Therefore, while features of p24 abundance or antibody titers may have an incomplete and complex relationship with progression, a p24/gp120 ratio may be more predictive.

Viremia was related to a broader set of components, most prominently 5 and (negatively) 6 (Fig. 6B). Given the importance of component 6 in predicting nearly all of the functional measurements (Fig. 6A), its importance in predicting viremia is to be expected. Component 5 likely serves to separate subjects based on their progression status as a proxy for antiretroviral therapy effect. That is, elite controllers, or those who control their infection without anti-retroviral therapy (ART), are likely to be immunologically distinct from treated progressors, who are not viremic due to ART. In all, a small number of consistent patterns accurately predict subjects’ HIV infection status. This is in stark contrast to previous approaches, where optimal models required 30–50 variables, and predictions rely on a complex collection of positive and negative associations with antigen and receptor measurements.^12^

## Discussion

We show here that structured data decomposition can improve our view of systems serology measurements. Specifically, this approach recognizes that measurements and their variation take place across distinct and separable dimensions—namely antigen binding and immune receptor engagement. Using this property, we identify that these measurements can be reduced more efficiently (Fig. 2), can be made robust to missing values (Fig. 3), and that properties of the immune system and infection can be accurately predicted (Fig. 5). Most critically, this form of dimensionality reduction provides a clearer interpretation of the resulting models (Fig. 4, 6), as it accounts for the high degree of inter-correlation across each dimension.

The key advancement of TMTF is separating the contribution of antigen from immune receptor binding. This separation enables the improved data reduction by avoiding repetition of the antigens for each receptor measurement or vice-versa in the factors (Fig. 2). Separation is also what allowed us to identify that just 8–10 consistent patterns exist within HIV subject’s serology (Fig. 4). Remarkably, even a multi-factorial change in the relationship between HIV and the immune response, like progression, could be accurately captured by a single pattern within the serology measurements (Fig. 4B, 6B). While each functional measurement was predicted through a combination of factors, component 6 contributed to nearly every prediction (Fig. 4B). This suggests that these functions are tuned through both shared and individualized regulatory changes. One limitation of the current immune functional assessments and glycan measurements is that they are not antigen-specific; future refinements to these measurements may reveal more precise regulation,^4^ particularly as glycans are known to be tuned in an antigen-specific manner.^32,33^

Advancements in the decomposition approach will continue to improve the predictions, interpretation, and robustness of systems serology data. While each antigen is treated similarly along one dimension, antigenic sites and strains could be separated into distinct dimensions before decomposition. This could lead to further data reduction (e.g. both strains of p24 and gp41 antigens share an identical signature; Fig. 4C), and simplify comparisons between strains. Interestingly, many of the subject components were highly correlated, indicating that components mostly separated due to differences in explaining the antigen and receptor dimensions (Fig. 4). As the subjects dimension results in the largest factor matrix (Fig. 4A), nested PCA or other factorization of it could easily reduce the data another two-fold. One current challenge is that some effect—possibly a technical artifact such as dilution differences—leads to non-linearities between receptors in the FcγR binding measurements (Fig. S2). Tensor generalizations of non-linear data reduction methods, like kernel-based factorization or autoencoders, could help to account for these patterns.^34^ To the extent that these different factors reflect differences in the avidity versus affinity of binding, factorization using a mechanistic binding model may help to separate these contributions.^35^ Conversely, TMTF is sure to have wider applications in other comprehensive molecular profiling experiments. Many datasets exist where measurements are made across varying dimensionality, such as with and without a temporal axis, and one wishes to find conserved patterns across the entirety of the data. As one example, TMTF could be used to look for patterns across paired time-course and endpoint single-cell measurements.^36^

More effective dimensionality reduction in turn enables new ways of viewing antibody-mediated protection. As mentioned above, systems serology in a way provides a view of antibody-mediated immunity akin to the microarray for gene expression data. The analogy reveals a few modeling advancements that will help leverage this data. One valuable property of TMTF is that it separates the immune receptor and antigen-binding patterns within the data. This will enable surveys for common Fc response patterns across diseases because these different datasets would still share this axis. This “transfer learning” could therefore help to identify common patterns of immune dysregulation. With more extensive profiling of the various glycosylation and isotype Fc forms, it would be possible to fix the receptor axis of the decomposition, to match new measurements to specific known immunologic patterns. These pattern-matching approaches would be much like gene set enrichment analysis for expression data.^37^ Finally, the binding interactions of antibodies, while they produce combinatorial complexity, are a simple set of antigen and receptor binding. Ultimately, one should be able to apply multivalent binding models to mechanistically model the interactions within serum.^35,38^ Such a mechanistic view would not only allow us to exactly identify mechanisms of antibody-mediated protection, but help to guide more advanced multi-modality therapeutic interventions, like monoclonal inhibitors or enhancers of antibody response that cooperate with the cocktail of endogenous antibodies.

Advancements outside systems serology will ultimately work alongside these measurements to expand our view of immunity. Much like how systems serology has served to profile antibody-mediated protection, profiling methods are helping to characterize T cell-mediated immunity.^39^ These technologies, alongside more traditional technologies to profile cytokine response, gene expression, and other molecular features, promise to provide a truly comprehensive view of immunity. Integrating these data will require dimensionality reduction techniques that recognize the structure of these data alone and in combination. Factorization methods will be a natural solution to this challenge, due to their scalability, “exibility, and interpretability.^20^

## Methods

All analysis was implemented in Python v3.8 and can be found at https://github.com/meyer-lab/systemsSerology.

### Subject cohort, antibody purification, effector function assays, and glycan analysis

All experimental measurements were used unmodified from prior work.^12^ The only exception is the gp140 antigen from the HVBc2 strain. This was found to be scaled much larger than the other antigens, and so was multiplied by 0.000001 to put it on a similar scale to other antigens. A small number of receptor binding measurements were found to have very large negative values. These were taken to be outliers and clipped to 0.0. HIV subjects were classified into four categories: untreated progressors, who failed to control viremia without combined anti-retroviral therapy (cART), treated progressors, who similarly failed to control viremia without cART but were on it for the study measurements, viremic controllers, who possessed a viral load between 50 and 2,000 RNA copies/mL without cART, and elite controllers, who had less than 50 copies/mL without cART. These were then grouped into two classifications: controllers (EC and VC) versus progressors (UP and TP); and viremic (UP and VC) versus non-viremic (TP and EC).

### Total Matrix-Tensor Factorization

We decomposed the systems serology measurements into a reduced series of Kruskal-formatted factors. Tensor operations were defined using Tensorly.^40^ To capture the structure of the data, where the majority of measurements were made for specific antigens, but gp120-associated antibody glycosylation was measured in an antigen-generic form, we separated these two types of data into a separate 3-mode tensor and matrix, with shared subject-dimension factors:

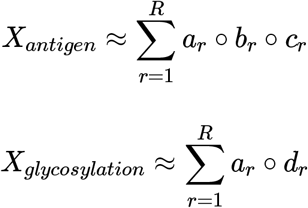

where *a*_*r*_, *b*_*r*_, and *c*_*r*_ are vectors indicating variation along the subject, receptor, and antigen dimensions, respectively. *d*_*r*_ is a vector indicating variation along glycan forms within the glycan matrix.

Decomposition was performed through an alternating least squares (ALS) scheme.^18^ Each least squares step was performed separately for each slice along a given mode, with missing values removed. While this made each iteration step much slower, convergence was much faster as a consequence of requiring fewer iterations. Missing values did not strictly follow a tensor slice pattern, and so alternative approaches such as a sampling Khatri-Rao product were disregarded as they would still require iterative filing.^41^ This strategy required many more iterations due to a high fraction of missing values (43%). The ALS iterations were repeated until the improvement in R2X over the last ten iterations was less than 1 × 10^−7^.

To enforce shared factors along the subject dimension, the antigen tensor and glycan matrix were concatenated after tensor unfolding. The Khatri-Rao product of the receptor and antigen factors was similarly concatenated to the glycan factors. The least-squares solution on this axis therefore solved for minimizing the squared error across both data compendiums. The other dimensions were solved using a standard ALS approach.

### Reconstruction Fidelity

To calculate the fidelity of our factorization results, we calculated the percent variance explained. First, the total variance was calculated by summing the variance in both the antigen-specific tensor and glycan matrix:

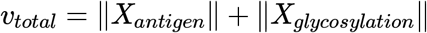

Any missing values were ignored in the variance calculation throughout. Then, the remaining variance after taking the difference between the original data and its reconstruction was calculated:

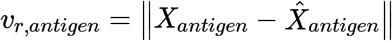

An analogous equation was used for the glycan matrix. Finally, the fraction of variance explained was calculated:

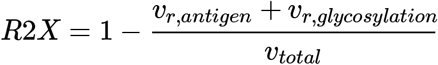

Where indicated, this quantity was calculated for values left out to assess the fidelity of imputation. In these cases this quantity was only calculated on those left out values, and indicated as Q2X.

### Cross-Validation

We employed a 10-fold cross-validation strategy to evaluate each prediction model. Cross-validation was stratified by class for classification. Unlike in earlier work,^12^ all subjects were randomly shuffled before cross-validation to prevent the influence of subject ordering in the dataset. We found that sharing the cross-validation fold structure between hyperparameter selection and model benchmarking led to consistent overfitting. For the factorization-based predictions, the better performing of either a linear or Gaussian process model was used. The linear model only showed superior performance when predicting ADCD.

### Logistic Regression / Elastic Net

Logistic regression and elastic net were performed using LogisticRegressionCV and ElasticNetCV implemented within scikit-learn .^42^ Both methods used 10-fold cross-validation to select the regularization strength with smallest cross-validation error, and a fraction of l1 regularization equal to 0.8 to match previous results.^12^ Logistic regression used the SAGA solver.^43^ Elastic net regression was set to normalize the data before model assembly.

### Gaussian Process Regression / Classification

Gaussian process regression or classification was performed where indicated using the implementation within scikit-learn .^42^ Regression was performed with output scaling. In both cases, an anisotropic radial basis function kernel was used with a constant scaling factor and additive white noise. The kernel’s parameters were left unbounded.

### Principal Components Analysis

Principal components analysis was performed using the implementation within the Python package statsmodels and the SVD-based solver. Missing values were handled by an expectation-maximization approach, wherein they were filled in with the imputed value. This filing step was performed up to 100 iterations until convergence as determined by a tolerance of 1 × 10^−5^.

### Missingness Imputation

To evaluate the ability of factorization to impute missing data, we introduced new missing values by removing chords from the antigen-specific tensor, and then looking at the variance explained on reconstruction (Q2X). More specifically, fifteen randomly selected receptor-antigen pairs were entirely removed and marked as missing across all subjects. TMTF decomposition was performed as described above, and then these left out data were compared to the reconstructed values. This process was repeated for the same chords across varying numbers of components.

## Acknowledgements

This work was supported by NIH U01-AI-148119 to A.S.M. The authors declare no competing financial interests. We thank Cyrillus Tan whose comments helped improve this manuscript.

## Author contributions statement

A.S.M. conceived of the study. All authors performed the computational analysis and wrote the paper.

## Supplementary Figures

**Figure S1:**
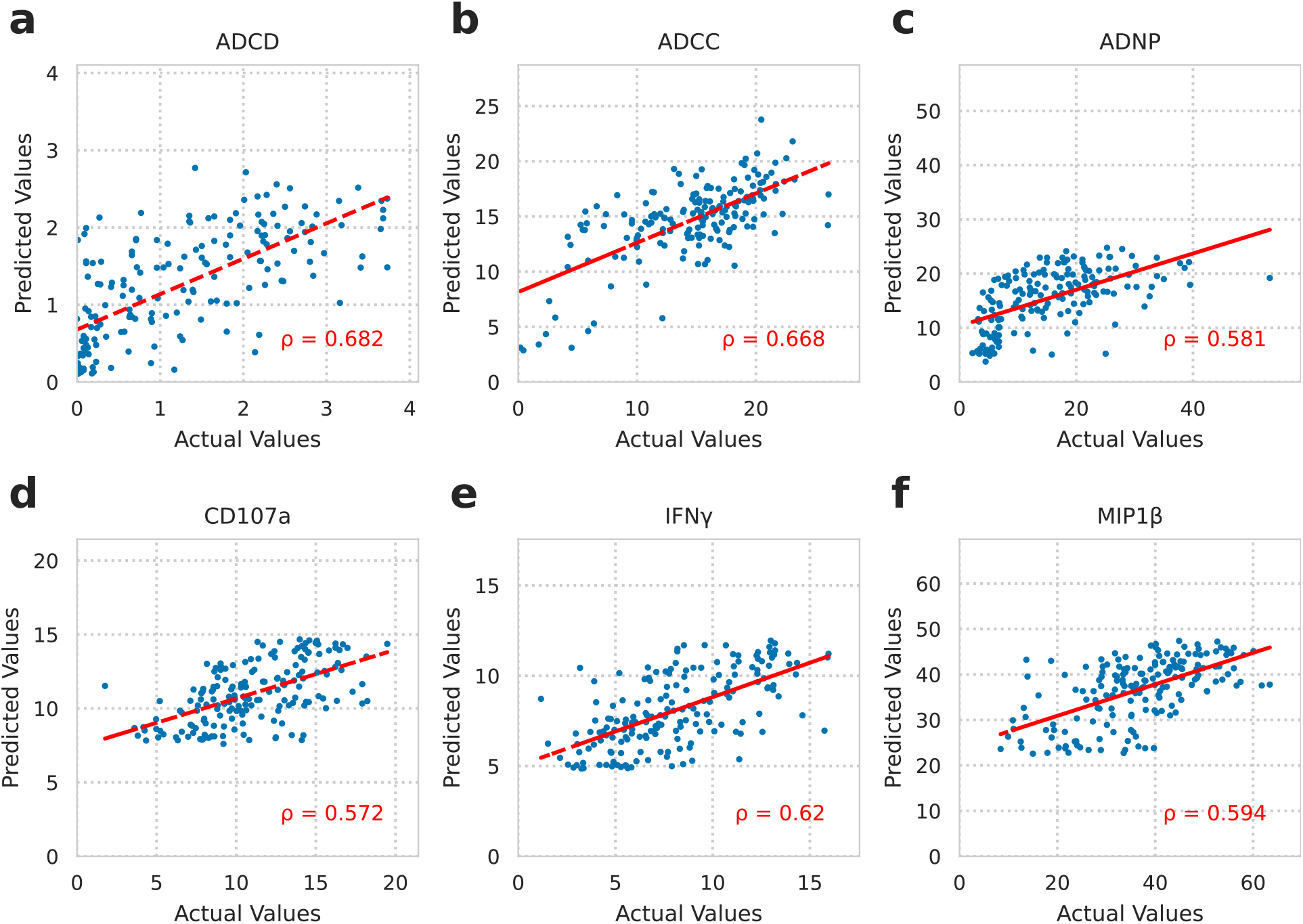
Predicted versus actual values of factorization-based regression models. Each point represents one subject and values represent antibody effector function assay values.^12^ Accuracy was calculated using the Pearson correlation coefficient. The red line represents the line of best fit.

**Figure S2:**
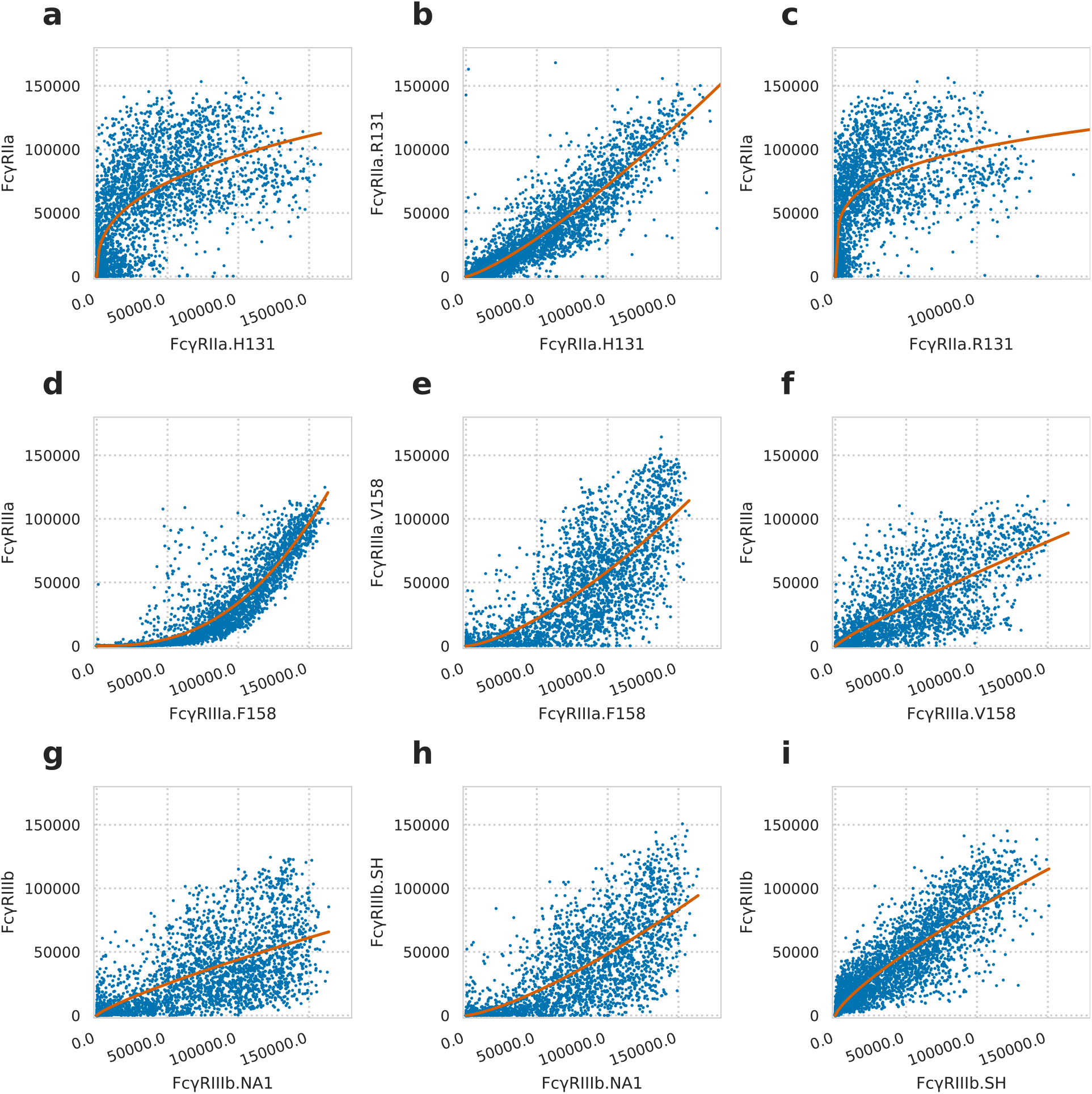
Comparing receptor genotype measurements across shared antigens shows consistent non-linear relationships. Plots comparing the binding among the FcγRIIa (a–c), FcγRIIIa (d–f), and FcγRIIIb (g–i) receptors, across all shared antigens and subjects. The red line shows the least-squares regression of a power curve. While one would expect sample-by- sample differences due to receptor affinity, the most prominent pattern is a consistent non-linear relationship. Also, the non-genotpe-specific receptor was the most divergent in measurement in every case. This suggests some consistent, technical artifact between the generic and genotype-resolved measurements.

